# Unpaired TCRα + TCRβ sequencing is sufficient for training machine learning TCR-epitope recognition predictors

**DOI:** 10.64898/2026.03.16.711991

**Authors:** Aisha Shah, Raphael Genolet, Aymeric Auger, Dana Léa Moreno, Yan Liu, Giancarlo Croce, Julien Racle, Alexandre Harari, David Gfeller

## Abstract

T-cell recognition of infected and malignant cells is elicited by the binding of heterodimeric T-Cell Receptors (TCRs) to epitopes and both the TCRα and the TCRβ chains play a key role in these interactions. Machine learning tools trained on databases of TCRs recognizing diverse epitopes are useful for identifying epitope-specific TCRs in large TCR repertoire datasets. However, collecting paired TCRαβ sequences to train such tools is associated with significant sequencing costs. Here we demonstrate that unpaired TCRα + TCRβ sequencing of epitope-specific T cells can be used for training TCR-epitope recognition predictors at a much reduced cost compared to standard single-cell TCR sequencing protocols and with no impact on prediction accuracy. Applying this approach to some unseen epitopes used in the IMMREP community benchmark demonstrates improved accuracy compared to both existing machine learning models and AlphaFold3-based predictions.

## Introduction

T-cell recognition of malignant or infected cells is elicited by the interactions between T-Cell Receptors (TCRs) and antigenic peptides presented on Major Histocompatibility Complex (MHC) molecules. TCRs are heterodimeric molecules with one α chain encoded at the TRA locus on chromosome 14 and one β chain encoded at the TRB locus on chromosome 7. Each TCR chain displays a very high sequence diversity as a result of the so-called V(D)J recombination. In this process, the α chain is formed by recombination of randomly selected Vα and Jα segments, and the β chain is formed by recombination of randomly selected Vβ and Jβ segments with a short Dβ segment between the Vβ and Jβ. Insertions and deletions further occur at the V(D)J junction, a region comprised within the Complementary Determining Region 3 (CDR3) loop.

Multiple studies have demonstrated that both chains play a central role in TCR-epitope recognition ^1–5^, with some epitopes showing more specificity in the α chain, like LLWNGPMAV restricted to HLA-A*02:01 (see https://tcrmotifatlas.unil.ch/model/A0201_LLWNGPMAV ^5^), and others displaying more specificity in the β chain, like TTDPSFLGRY restricted to HLA-A*01:01 (see https://tcrmotifatlas.unil.ch/model/A0101_TTDPSFLGRY). Besides the specificity encoded in each chain, some specificity has been reported in the pairing of the chains ^1,6^. For instance, not all combinations of α and β chains derived from TCRs binding to an epitope preserve the binding to this epitope.

Experimental identification of TCRs recognizing a given epitope can be done by TCR-sequencing of epitope-specific T cells. Typically, cells are stimulated with a peptide and put in culture for a few days in a medium that favors expansion of epitope-specific T cells. T cells with epitope-specific TCRs can be sorted with peptide-MHC multimers or antibodies recognizing T-cell activation markers. The TCRs of the sorted T cells are then sequenced. Two main categories of TCR-sequencing technologies exist. In the first category, specific primers are used to amplify separately the TCRα and TCRβ loci, or only the TCRβ locus in some popular protocols like ImmunoSeq, and bulk sequencing is applied ^7–9^. This approach reaches a high sequencing depth and can analyze millions of cells per sample. The cost for library preparation + TCR-sequencing ranges from $300 to $2000 per sample with commercial technologies from Takara, iRepertoire or Adaptive Biotechnologies, or when using manually designed primers for PCR amplification ^7,9–11^. However, it does not provide information about which TCRα chain was in complex with which TCRβ chain, and in silico pairing based on frequencies can be done accurately only for a small minority of clones ^12,13^. In the second category, the full TCRαβ sequences are determined for each T cell. Many approaches rely on single-cell sequencing techniques, where each T cell is encapsulated in a droplet and TCRα and TCRβ reads in each droplet are labelled with the same barcode ^14,15^. Recent technological developments by companies like Parse Biosciences, Omniscope have increased the throughput of these approaches. Alternatively, studies have shown how to recover actual TCRαβ pairs from unpaired TCRα + TCRβ sequencing after distributing a sample into multiple wells and computationally pairing co-occurring TCRα and TCRβ chains ^16,17^. Many of these paired TCRαβ sequencing pipelines have lower throughput and higher costs (e.g., ∼$2000 for a few thousand cells with the popular 10x Genomics toolkit) than bulk TCRα + TCRβ sequencing, or require dedicated setup which are not available in every lab.

Many machine-learning tools have been developed to predict TCR-epitope interactions by learning patterns from epitope-specific TCR sequences ^1,2,5,18–22^. Several of these tools can score millions of TCRs in a few minutes, making them attractive to annotate large TCR repertoires from patient cohorts for multiple epitopes. Recent benchmarks have demonstrated that reasonable predictive power can be reached for epitopes with abundant and reliable training data, but performance drops substantially for epitopes with only a few known TCRs and is close to random for epitopes with no previously characterized TCRs ^23,24^. The latter are commonly referred to as ‘unseen’ epitopes. These tools reach the best accuracy when considering both the TCRα and TCRβ chains, which is consistent with 3D structures where both chains make on average the same number of contacts with the epitope ^4,5^. For this reason, several recent tools focus on paired TCRαβ sequences of epitope-specific T cells in their training ^20,21,25–28^. However, it remains unclear how much information related to the pairing itself (i.e., after regressing out the specificity encoded in each chain) contributes to TCR-epitope prediction accuracy. Approaches using the latest protein structure and interaction modelling tools ^29–31^ show promise for unseen epitopes, although their predictive power varies across different epitopes ^32–34^.

Here we demonstrate that machine learning TCR-epitope recognition prediction tools can be trained on unpaired TCRα + TCRβ sequencing data from epitope-specific T cells and illustrate how this approach could be applied to previously unseen epitopes used in the IMMREP23 benchmark ^23^.

## Results

### Shuffling chains in paired TCRαβ sequences used for training TCR-epitope interaction predictors preserves their accuracy

To explore the use of unpaired TCRα + TCRβ sequences for training predictors of TCR-epitope interactions, we used a collection of publicly available paired TCRαβ sequences known to interact with multiple epitopes ^5,35,36^. For each epitope with at least 10 complete TCRαβ sequences, we performed a five-fold cross-validation and compared model performance when training on either actual paired TCRαβ sequences or after randomly shuffling α and β chains in paired TCRαβ sequences (see Materials and Methods, Table S1 and Figure 1A). The shuffling of chains ensures that all the specificity encoded in each chain is kept, while the potential specificity encoded in the pairing between chains is removed. This analysis was done for three state-of-the-art tools that can be retrained: MixTCRpred ^20^, NetTCR2.2 ^21^ and TULIP^22^. Across all three tools, models trained on paired or shuffled TCRαβ data achieved highly similar predictive performance, as measured by normalized Area Under the receiver operating Curve up to a false-positive rate of 0.1 (AUC01) (Figure 1B, Figure S1A). This finding was robust to alternative evaluation choices, like using AUC instead of AUC01 (Figure S1B-C) or evaluating on test sets having swapped negatives derived from TCRs binding other epitopes instead of TCRs with undetermined specificities (Figure S1D). To exclude the possibility that our observations would result from the presence of epitopes with relatively low number of TCRs or with limited specificity in the data which could prevent learning α-β correlations, we restricted the analysis to epitopes with at least 200 paired TCRαβ sequences or to epitopes with AUC0.1 >= 0.8 when trained on paired data. Here again, no difference could be observed between predictions trained on paired TCRαβ or shuffled TCRαβ sequences (Figure S1E-F).

**Figure 1:**
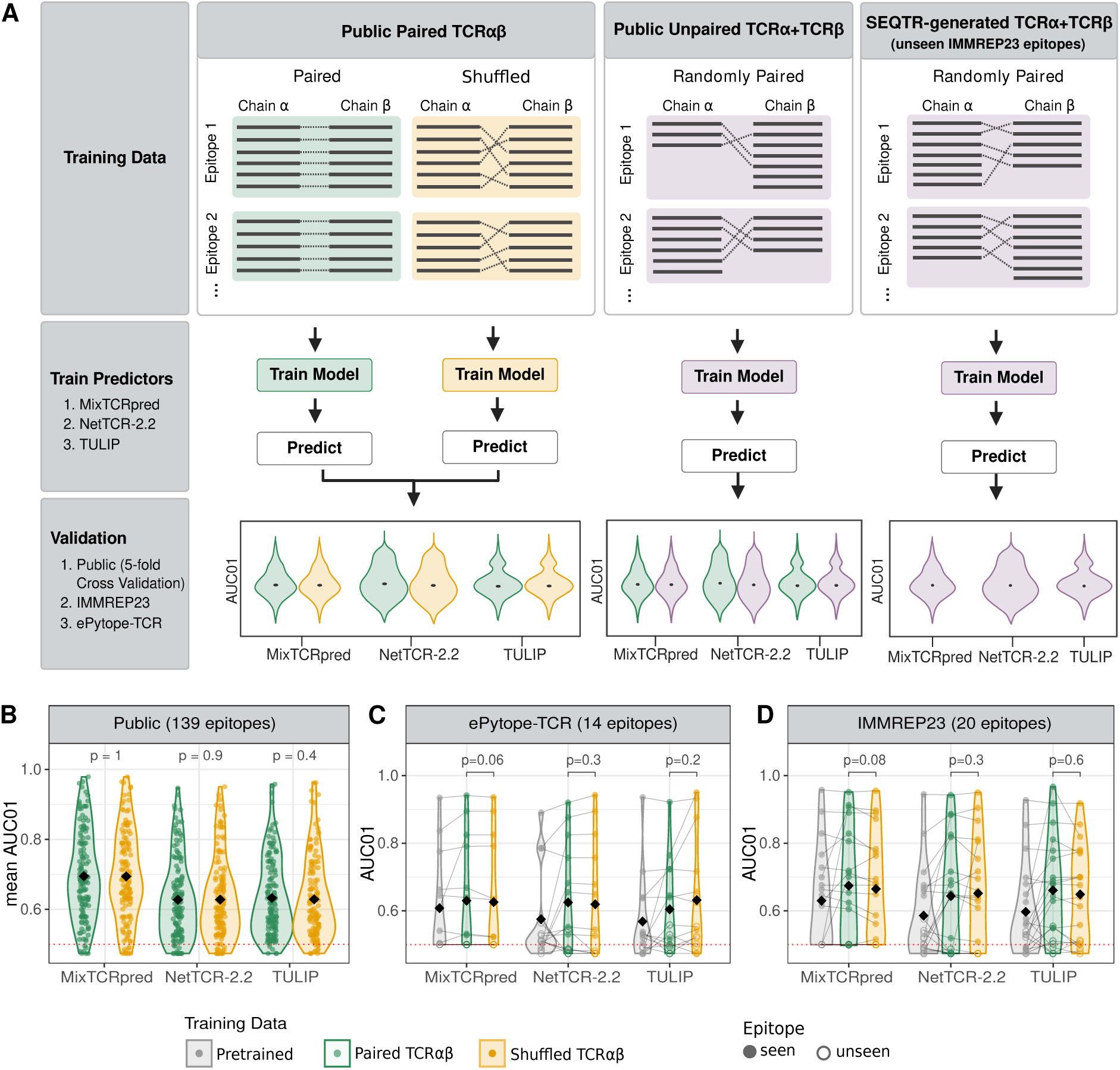
Shuffling chains in paired TCRαβ sequences used for training TCR-epitope interaction predictors preserves their accuracy. **A)** Overview of the different scenarios considered in this study to compare TCR-epitope interaction prediction tools trained on paired TCRαβ, on shuffled TCRαβ or on public or newly generated unpaired TCRα + TCRβ sequences. **B)** Cross-validation AUC01 values obtained when training different tools on paired TCRαβ or shuffled TCRαβ sequences (see Figure S1A for the distribution of the differences in AUC01 values). **C)** AUC01 values for epitopes in the ePytope-TCR benchmark dataset when training different tools on paired TCRαβ or shuffled TCRαβ sequences. **D)** AUC01 values for epitopes in the IMMREP23 benchmark dataset when training different tools on paired TCRαβ or shuffled TCRαβ sequences. P-values in panels B, C and D were computed with the Wilcoxon paired test. Black diamonds show the mean values across all epitopes. For comparison, AUC01 values obtained with the published models of each tool are shown in grey in panels C and D.

To further explore the impact of the TCRαβ chain pairing, we used the ePytope-TCR ^24^ and IMMREP23 ^23^ external benchmark datasets. For each epitope in these datasets the different tools were trained on the same set of publicly available data, excluding any data that was generated for these two competitions (see Materials and Methods). As before, we could not observe consistent differences in AUC01 or AUC values when training tools on paired TCRαβ or shuffled TCRαβ data (Figure 1C-D, Figure S1G-H).

Overall, these results indicate that potential information encoded in the pairing between TCRα and TCRβ chains does not contribute much to the accuracy of current TCR-epitope interaction predictions tools, including for epitopes with large training data (i.e., >200 TCRs) and/or strong specificity signals in the data (i.e., AUC01>=0.8).

### Unpaired TCRα + TCRβ sequences can be used for training TCR-epitope interaction predictors

We next explored whether unpaired TCRα + TCRβ sequencing data can be used to train predictors of TCR-epitope interactions without compromising their performance. We first focused on publicly available data and selected epitopes for which at least 10 paired TCRαβ, 10 single-chain TCRα and 10 single-chain TCRβ sequences were available, resulting in a total of 24 epitopes. For each epitope, we compared model performance when training on paired TCRαβ sequences versus randomly paired TCRα + TCRβ sequences, ensuring balanced size in the training data for both models for each epitope (Figure 1A) (see Materials and Methods and Table S2). Despite potential differences in quality between paired and unpaired data, no statistically significant difference in AUC01 values was observed between models trained on paired and unpaired TCRs (Figure 2A, see Figure S2A for AUC values). We extended this analysis to epitopes in the ePytope-TCR and IMMPREP23 benchmark datasets. Consistent with our previous observations, training on unpaired TCRα + TCRβ sequences yielded AUC01 values that were similar to those obtained using paired TCRαβ data for most epitopes (Figure 2B-C and Figure S2B-C for AUC values).

**Figure 2:**
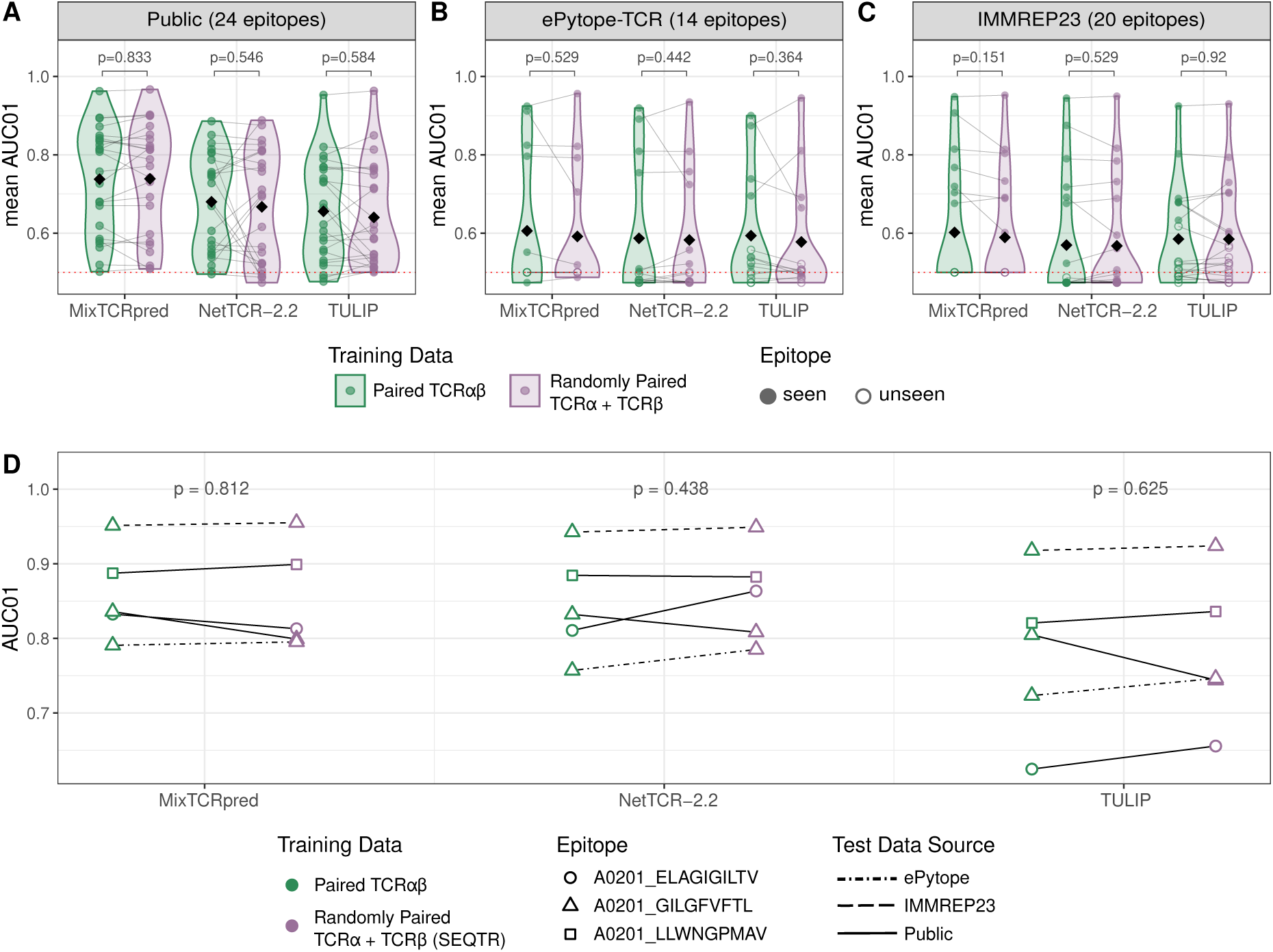
Unpaired TCRα + TCRβ sequences can be used for training TCR-epitope interaction predictors. **A)** AUC01 values obtained when training tools on paired TCRαβ or unpaired TCRα + TCRβ data and benchmarking them using the same cross-validation setting as in Figure 1B for the subset of epitopes with both paired and unpaired training data. **B)** AUC01 values across ePytope-TCR epitopes when training tools on paired TCRαβ or unpaired TCRα + TCRβ data. **C)** AUC01 values across IMMREP23 epitopes when training tools on paired TCRαβ or unpaired TCRα + TCRβ data. **D)** AUC01 values for three epitopes (GILGFVFTL, LLWNGPMAV and ELAGIGILTV, all restricted to HLA-A*02:01) when training tools on paired TCRαβ or unpaired TCRα + TCRβ data generated with the SEQTR protocol. GILGFVFTL predictions were evaluated on three test sets indicated by different line types. P-values were computed with the Wilcoxon paired test. Black diamonds show the mean over all epitopes.

To further assess the practical use of generating unpaired TCRα + TCRβ sequencing for training predictors, we used an independent experimental dataset of epitope-specific unpaired TCRα + TCRβ sequences recently determined in our group ^5^. Briefly, T cells from HLA-A*02:01 positive donors had been stimulated with three well-characterized HLA-A*02:01 restricted viral and tumor-derived peptides: GILGFVFTL from influenza, LLWNGPMAV from yellow fever and ELAGIGILTV from the Melan-A cancer antigen ^5^. Epitope-specific T cells had been sorted with the corresponding peptide-MHC multimers and their TCRα and TCRβ repertoires had been sequenced with SEQTR ^9^. We compared the performance of MixTCRpred, NetTCR, and TULIP trained either on publicly available paired TCRαβ data or on the newly generated unpaired TCRα and TCRβ sequences for the three epitopes in the previously considered benchmarking datasets where they appeared (i.e., cross-validation for all three epitopes, plus IMMREP23 and ePytope-TCR for A0201_GILGFVFTL). Here as well, the predictive power was very similar across the two training strategies (Figure 2D and Figure S2D for AUC values).

Altogether, these results demonstrate that unpaired TCRα + TCRβ sequencing data can be used to train predictors of TCR-epitope recognition without compromising accuracy compared to models trained on paired TCRαβ data.

### Unpaired TCRα + TCRβ sequencing with SEQTR enables predictions for previously unseen epitopes

To demonstrate how unpaired TCRα + TCRβ sequencing can be used to train predictors for epitopes without known TCRs, we selected three epitopes (i.e., A0101_SALPTNADLY, A0101_TDLGQNLLY and A0101_VSDGGPNLY) used in the IMMREP23 competition with very little (12 TCRβ chains for A0101_TDLGQNLLY and 3 TCRαβ for A0101_VSDGGPNLY) or no (A0101_SALPTNADLY) training data available in VDJdb ^37^ at the time of the IMMPEP23 competition. For each epitope, we collected T cells from HLA-A*01:01 positive donors and stimulated them with the corresponding peptides (Figure 3A). After stimulation with each peptide for 10 days, the T cells were sorted based on CD137 upregulation (Figure S3) and the TCRα and TCRβ repertoires were sequenced with the SEQTR protocol ^9^. This led to the identification of hundreds of epitope-specific TCRα and TCRβ chains (Table S3). As indicated in the IMMPEP23 publication ^23^, these experiments were performed within the one month timeframe of the competition. The cost of library preparation and TCR sequencing with SEQTR was $350 per sample, which makes this approach scalable and cost-efficient.

**Figure 3:**
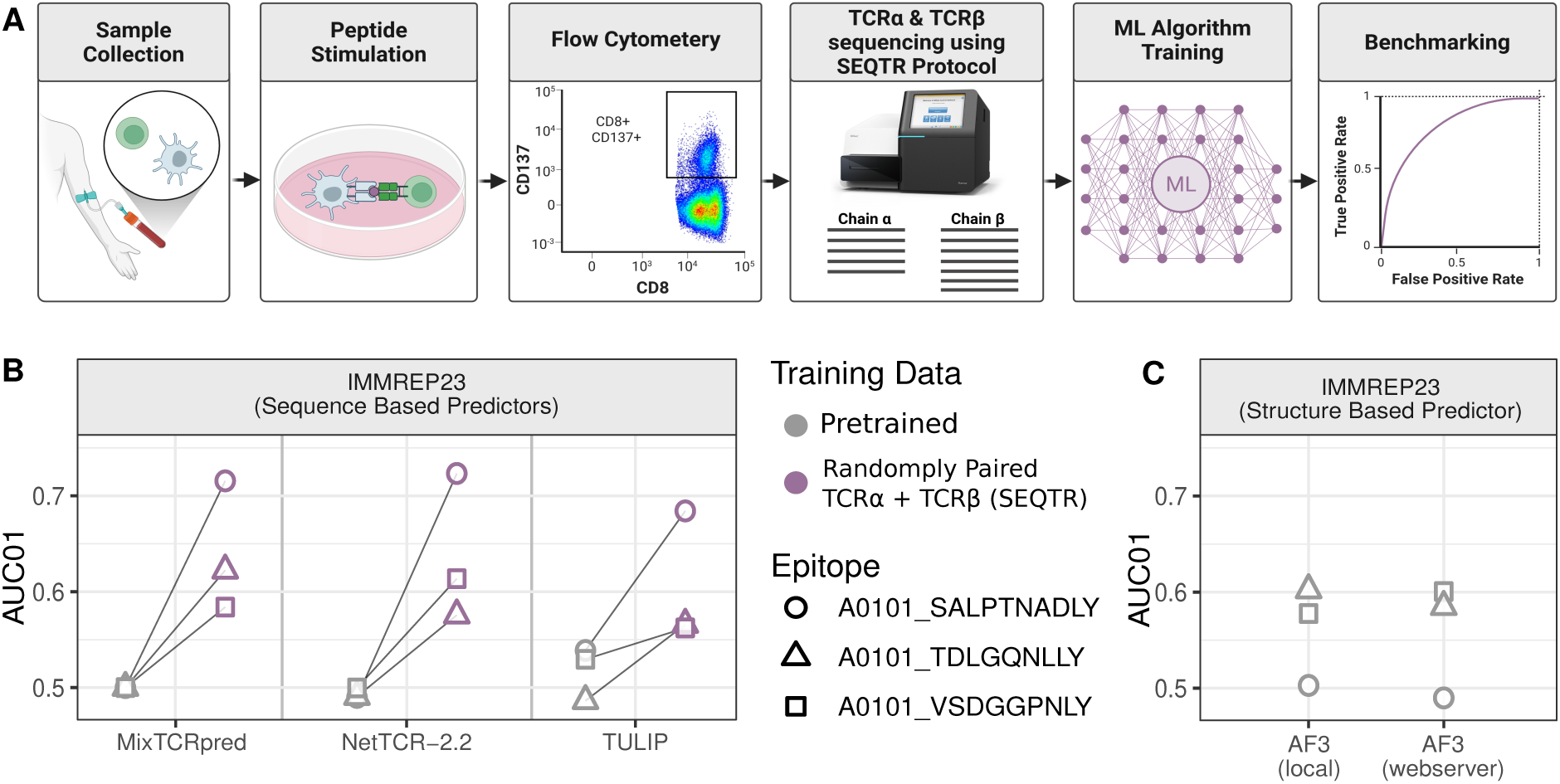
Unpaired TCRα + TCRβ sequencing enables predictions for previously unseen epitopes. **A)** Illustration of the proposed pipeline to generate training data for unseen epitopes based on TCRα + TCRβ sequencing with SEQTR. **B)** Mean AUC01 obtained for tools trained with or without the newly generated unpaired TCRα + TCRβ sequencing data for three IMMPEP23 epitopes with very limited or no publicly available training data. **C)** AUC01 obtained with the ipTM of AF3 for the three IMMPEP23 epitopes considered in panel B.

We then trained MixTCRpred, NetTCR and TULIP with the randomly paired TCRα + TCRβ SEQTR data using 5 different seeds for random pairing (Table S4). Model performance was evaluated on the IMMREP23 benchmark dataset. For all three epitopes the models trained on SEQTR data displayed clear, although not perfect, predictive power (Figure 3B), while no predictive power was observed when using published pretrained models which did not include the new data in their training.

To explore the molecular basis of these predictions, we compared the TCR specificity profiles ^5^ of the SEQTR data to those of the IMMREP23 data (Figure S4). For all epitopes, the IMMREP23 and SEQTR datasets showed some similarity in V and J gene usage and CDR3 length distributions. These include for instance enrichment for TRAV1-2, TRAV12-3, TRBV2 and TRBJ2-7 in TCRs recognizing the A0101_SALPTNADLY epitope, for which models trained on the SEQTR data had the best predictive power.

We further compared these predictions with structure-based modeling using AlphaFold3 (AF3), either via a local command-line implementation or via the webserver (Figure 3C). AUC01 values were calculated based on the interface predicted template modelling (ipTM) scores computed by AF3 between the TCR (i.e., TCRα and TCRβ) and the epitope (i.e., peptide and MHC) chains (see Materials and Methods). For A0101_SALPTNADLY, all sequence-based predictors trained on SEQTR data consistently demonstrated much better predictions than AF3, while for the two other epitopes prediction accuracy was either similar for MixTCRpred and NetTCR and slightly lower for TULIP.

Overall, these results demonstrate that unpaired TCRα + TCRβ sequencing can be used to train predictors of TCR-epitope recognition for previously unseen epitopes, following the pipeline in Figure 3A. This highlights the practical utility of cost-efficient unpaired TCRα + TCRβ sequencing for profiling epitope-specific TCRs and enhancing the epitope coverage of machine learning TCR-epitope recognition predictors.

## Discussion

Accurate TCR-epitope recognition predictions could have a major impact for both fundamental understanding of immune responses and clinical applications in infectious diseases, autoimmunity and cancer immunotherapy. As both the TCRα and the TCRβ chains play a central role in TCR-epitope recognition, paired TCRαβ data have been used to train many recent state-of-the-art tools. However, single-cell technologies remain comparatively expensive to generate such data and yield lower sequencing depth than approaches based on unpaired TCRα + TCRβ profiling. Recent methodological advances that have massively improved the recovery of paired TCRαβ sequences in bulk TCR repertoires ^17^ are also more challenging to apply to the relatively small number of epitope-specific T-cells typically obtained from biological samples. In this work, we demonstrate that unpaired TCRα + TCRβ sequencing of epitope-specific T cells is sufficient for training machine learning TCR-epitope recognition predictors, thereby allowing performing such experiments at significantly reduced costs.

Across multiple epitopes, benchmark datasets and prediction tools, we consistently observed that preserving the actual TCRαβ pairing in the training data had little to no impact on predictive performance, although MixTCRpred, NetTCR and TULIP could all in theory learn specificity encoded in the pairing between chains. Several factors likely contribute to this observation. First, for some epitopes, close to no specificity is encoded in the pairing between α and β chains. This is typically the case when a strongly dominant V/J gene usage is found in both chains ^1,5^. In such cases, the key information is captured at the level of individual chains, leaving little additional signal to be gained from modeling TCRαβ pairing. Second, for many epitopes, the number of training examples required to learn potential correlations between chains is simply too low in today’s data. It should be noted that learning correlation signals in combinations of features, such as specific Vα-Vβ pairs showing higher joint frequency than what is expected from their individual point frequencies, requires much more data than for individual features, such as specific Vα and specific Vβ usage. Third, we cannot exclude that all existing tools may have failed at learning some of the correlations, even in cases where enough information would be available in today’s data. Finally, it is important to realize that TCRαβ chain pairs that do not bind to an epitope despite having a good TCRα and a good TCRβ chain represent only a very tiny fraction of a TCR repertoire ^5^. Most of them will also not be selected and amplified in vivo, since they do not recognize an epitope. Therefore, in realistic scenarios where the task is to find epitope-specific TCRs from ∼10’000 TCRs sequenced in a patient, the probability of having even one of these non-binding pairs is quite low.

These observations indicate that correlations between α and β chains, even when they exist ^6^, are not critical to learn for many practical applications of TCR-epitope recognition predictions. This result is fully consistent with our recent observation that a predictor treating separately each chain reaches similar or better accuracy compared to other tools that can in theory learn complex correlation patterns across chains ^5^.

When training tools on unpaired TCRα + TCRβ sequences, we randomly paired them without duplicating chains and therefore discarded some of the TCRs in the chain with most data. This was done since many tools ^20,21^ used in our benchmark require paired TCRαβ in their input. An alternative approach could consist in developing machine learning algorithms that naturally support unpaired TCRα + TCRβ sequences in their training data ^5,22^.

The question whether TCR-epitope prediction tools can be trained on single-chain TCRs, like TCRβ only, has been addressed in previous studies ^4,5^. These studies have demonstrated that information from both chains is critical to reach the best predictive power. Combined with the results obtained in this study, this strongly supports the idea that retrieving the TCRα and TCRβ chains in epitope-specific T cells is critical but information about the exact pairing is not necessary when using these data to train predictors of TCR-epitope recognition.

Overall, our results demonstrate that unpaired TCRα + TCRβ sequencing of epitope-specific T cells captures sufficient information to train predictors of TCR-epitope recognition for many practical applications of such tools. The lower costs and enhanced sequencing depth make this approach promising to expand the coverage of clinically relevant epitopes for which machine learning TCR-epitope interaction predictions can be performed.

## Materials and Methods

### Data collection

Human TCRs recognizing multiple epitopes were retrieved from Liu et al. ^5^. These include data from VDJdb ^35^ and IEDB ^36^, as well as specific studies manually added to this collection. Both paired TCRαβ as well as single-chain TCRα or TCRβ were considered. Data from the 10X Genomics study ^38^ were excluded due to the reported high level of contamination ^20^. Data from the McPAS-TCR database ^39^ were not considered due to the reported issues with V/J names ^40^. In total, 139 epitopes had at least 10 paired human TCRαβ. Among them 24 also had at least 10 single-chain TCRα and 10 single-chain TCRβ sequences. Data from the ePytope-TCR ^24^ and IMMREP23 ^23^ benchmark were used as published in the original studies.

### Training tools on publicly available paired TCRαβ

All prediction tools (i.e., MixTCRpred (v1.0) ^20^, NetTCR (v2.2) ^21^ and TULIP (v1.0) ^22^) were trained using paired TCRαβ. NetTCR was used in the epitope-specific mode. This configuration is also the only mode available for MixTCRpred. To mimic realistic applications of TCR-epitope interaction predictions and avoid emergence of specificity patterns in negatives, non-binding TCRs needed for training MixTCRpred and NetTCR were retrieved from TCR repertoires with undetermined specificities in a fraction of ten negatives for each positive ^5^. TULIP was always trained on the full dataset of paired TCRαβ of multiple epitopes (by construction TULIP does not explicitly use negatives but requires TCRs binding to multiple epitopes for its training). Exact TCR sequences for each epitope and each cross-validation instance can be retrieved from Table S1.

### Training on shuffled paired TCRαβs

To assess the contribution of native TCRαβ pairing to predictive performance, we generated shuffled TCRαβ datasets in which TCRα and TCRβ chains were randomly reassigned within each epitope (Figure 1A). This procedure disrupts the original biological α-β pairing while preserving the specificity of individual α and β chains. All models were retrained using these shuffled TCRαβ combinations. Negatives in the training data remained the same as those used for the models trained on original paired TCRαβ data. The exact shuffled TCRαβ combinations generated for each epitope and cross-validation split are provided in Table S1.

### Training on publicly available unpaired TCRα + TCRβ

In this setting, MixTCRpred, NetTCR, and TULIP were trained using publicly available unpaired TCRα + TCRβ sequences. For each epitope, duplicated TCRα and TCRβ sequences were removed and the chain with the highest number of sequences was subsampled to have the same number of sequences for both chains. The remaining TCRα and TCRβ sequences were randomly paired to create the training datasets. Negatives in training data required for MixTCRpred and NetTCR were collected from TCR repertoires with undetermined specificities, in a fraction of ten negatives for each positive. The full list of randomly matched TCR sequences for each epitope can be retrieved from Table S2.

### Evaluating models trained on public data

#### Cross-validation on publicly available data

For models trained on public paired TCRαβ or shuffled TCRαβ, performance was assessed using the 5-fold cross-validation framework of ref. Liu *et al* ^5^. For each epitope with at least 10 TCRαβ sequences, we split these sequences into five groups. One group was used for testing, and the other four groups were used for training. To prevent information leakage between training and test data, any TCR in the cross-validation training set sharing the same CDR3α or CDR3β sequence as a TCR in the corresponding test set was excluded from the training data. To avoid issues with batch effects shared between negatives used in the training and the test set, negatives in the test were selected from studies not used to generate the negatives in the training set. For Figure S1D, swapped negatives were generated by randomly selecting TCRs binding to other epitopes instead of TCRs with undetermined specificity ^5^. For each epitope, the mean AUC01 or AUC across all five cross-validation sets was used.

When comparing models trained on public paired TCRαβ sequences and those trained on public unpaired TCRα + TCRβ sequences, one model per epitope was trained on unpaired TCRα + TCRβ data and evaluated on each of the five test sets from the cross-validation framework mentioned above. Reported performance corresponds to the mean AUC01 across the five test sets. For this comparison, the size of the training sets for models trained on paired and unpaired data were made equal by random subsampling. This was done to ensure that differences in predictive power would not result from different sizes of the training sets, since for some epitopes, paired TCRαβ data were much more abundant than unpaired TCRα + TCRβ, while the opposite was true for other epitopes.

#### External evaluation on IMMREP23 and ePytope-TCR datasets

External validation was performed using the ePytope-TCR (14 epitopes) and IMMREP23 (20 epitopes) benchmark datasets ^23,24^. Each benchmark consists of epitope-specific TCRs (positives) together with TCRs known to bind other epitopes considered in these benchmarks (‘swapped’ negatives). These datasets were used as independent test sets to assess the performance of models trained under different training configurations, namely paired TCRαβ, shuffled TCRαβ, and unpaired TCRα + TCRβ data, as well as pretrained models. To prevent data leakage, all TCRs originating from the IMMREP23 benchmark or from the ImmunoScape study ^41^ were excluded from the training sets of any model evaluated on the IMMREP23 benchmark. When comparing models trained on public paired TCRαβ sequences and those trained on public unpaired TCRα + TCRβ sequences, the size of the training sets was made equal by random subsampling.

When comparing models trained on paired and shuffled TCRαβ, sufficient data (>= 10 TCRs) was available for 14 of 20 IMMREP23 and 8 of 14 ePytope-TCR epitopes. When comparing models trained on paired TCRαβ versus unpaired TCRα + TCRβ, 7 IMMREP23 and 6 ePytope-TCR epitopes had both sufficient paired (>= 10 paired TCRαβ sequences) and unpaired data (>= 10 TCRs for each chain). For these epitopes with sufficient data, epitope-specific models were trained for MixTCRpred and NetTCR-2.2 and evaluated using AUC01. For epitopes lacking sufficient data, MixTCRpred AUC01 was set to 0.5, and NetTCR-2.2 predictions were made using pan-specific models trained on the remaining epitopes in the corresponding benchmark. For all analyses, TULIP predictions were generated using pan-epitope models trained on all the epitopes belonging to the respective benchmark.

For comparison, we also reported the AUC01 obtained with the pretrained version of each tool as published in the original study with no retraining (grey boxplots in Figure 1C-D). For MixTCRpred, pretrained models available for 11 IMMREP23 and 8 ePytope-TCR epitopes were downloaded from https://github.com/GfellerLab/MixTCRpred/tree/main (commit id: 950313a). For remaining epitopes AUC01 was set to 0.5. For NetTCR-2.2 pretrained epitope-specific models were available for 7 IMMREP23 and 4 ePytope-TCR epitopes. For remaining epitopes, a pretrained pan model was used. These epitopes are considered unseen epitopes by this published pan model. All these NetTCR-2.2 models (version t.1.v.2) were downloaded from https://github.com/mnielLab/NetTCR-2.2/tree/main (commit id: 244bb88). For TULIP, the pretrained model was downloaded from https://github.com/barthelemymp/TULIP-TCR/tree/main (commit id: 798fab9). 11 of ePytope epitopes and 16 IMMREP23 epitopes were present in training data of this published model of TULIP while remaining epitopes were considered unseen.

### Collection of T cells

Peripheral blood was obtained from donors enrolled under protocols approved by the Ethics Committee from Lausanne University Hospital, Switzerland (2017-00490) and all patients gave written informed consent.

#### In vitro stimulation of peripheral blood mononuclear cells (PBMC) with peptides

PBMCs were thawed, counted and rested overnight in RPMI (Gibco) supplemented with 10% FBS (Gibco) and 1% penicillin/streptomycin (BioConcept). After resting, PBMC were counted, washed with PBS and resuspended at 10e6 cells / mL in RPMI supplemented with 8% HuAB serum (Biowest), 1% penicillin/streptomycin and 100 U/mL of IL-2 (Proleukin, Novartis Pharma). 2 mL of cell suspension were plated per well of a 24 well plate and peptides were added at a final concentration of 1µM. Cells were incubated at 37°C for 10 days and, every 2-3 days, half of the cell culture media was replaced by fresh media. Cells were also split when confluence was reached.

#### Sorting of peptide-reactive CD8 T cells by CD137 upregulation

10 days after the initiation of the IVS, cells were re-challenged with peptides at a concentration of 1µM. 5x10e5 cells were kept unstimulated as background control. After an overnight incubation at 37°C, cells were harvested, washed with PBS and stained with a mix of the following reagents: Far Red dead cells stain (Invitrogen ref. L10120), anti-human CD4 Pacific Blue (BD Biosciences ref. 558116), anti-human CD8 FITC (BioLegend ref. 344704) and anti-human CD137 PE (Miltenyi Biotec ref. 130-119-885). After 20-30 minutes of incubation at 4°C, cells were washed and resuspended in PBS and sorting of the live CD4 - CD8 + CD137 + fraction was performed using a BD FACS Melody (BD Biosciences). Sorted cells were washed with PBS, resuspended in 300 µL of lysis/binding buffer (Invitrogen ref. A33562) and frozen at -80°C until TCR sequencing.

### TCR-sequencing and reconstruction

TCR sequencing of sorted CD8 T cells was performed with the SEQTR pipeline for both TCRα and TCRβ chains ^9^. Raw sequencing data were processed with MiXCR (Version: 4.7.0) to reconstruct TCRα and TCRβ chains ^42^ using the *generic-amplicon-with-umi.yaml* (https://github.com/milaboratory/mixcr/blob/476e4ad3b2457c9413a1cf05ae67a668d8df05b0/regression/presets/analyze/generic-amplicon-with-umi.yaml#L4) preset adapted to the SEQTR protocol. Specifically, align.species was set to “hsa”, tagPattern was defined according to the SEQTR primer design, tagMaxBudget was set to 15, align.parameters.vParameters.geneFeatureToAlign was set to “VTranscriptWithP”, align.parameters.vParameters.parameters.floatingLeftBound was set to false, and align.parameters.cParameters.parameters.floatingRightBound was set to true. In addition, assemble.cloneAssemblerParameters.separateByV, assemble.cloneAssemblerParameters.separateByJ, exportClones.filterOutOfFrames, and exportClones.filterStops were set to true.

The resulting dataset was then preprocessed using a series of quality control steps. A minimum threshold of five unique molecular identifiers was applied to keep only high-confidence sequences. Allele information was removed from V and J gene annotations and gene names were standardized to IMGT references. CDR3 sequences were validated by checking that the first amino acid and the final two amino acids matched the expected germline V and J gene sequences. CDR3 sequences shorter than 7 amino acids or longer than 22 amino acids were not considered.

### Training models on SEQTR-derived TCR data

For publicly available SEQTR datasets (A0201_GILGFVFTL, A0201_LLWNGPMAV, and A0201_ELAGIGILTV), no additional preprocessing was performed, as the data had already been processed in the original study ^5^. TCRα and TCRβ sequences were randomly paired by subsampling the chain with the largest number of sequences for each epitope, and the resulting randomly paired chains were used to retrain the different models. For newly generated data corresponding to IMMREP23 epitopes with very few or no training data available in public datasets (i.e., A0101_SALPTNADLY, A0101_TDLGQNLLY and A0101_VSDGGPNLY), TCRα and TCRβ sequences were randomly paired using five different random seeds. For MixTCRpred and NetTCR, negatives in the training sets were selected from repertoires of T cells with undetermined specificity and sequenced with the SEQTR protocol ^9^. The same set of negatives was used for all five randomly paired datasets, ensuring that only the α-β pairing of binding TCRs differed across seeds. Epitope-specific models were trained for MixTCRpred and NetTCR. For TULIP, a pan-epitope model was trained using the SEQTR data for all epitopes with such data, as well as data from 24 other epitopes used in training models on randomly paired TCRα + TCRβ sequences from public data.

### Evaluating models trained on SEQTR data

For A0201_GILGFVFTL, A0201_LLWNGPMAV, and A0201_ELAGIGILTV for which abundant data exist in public databases, we evaluated SEQTR-trained models on each of five cross-validation test sets and computed the mean AUC and AUC01. Performance was then compared to models trained on publicly available paired TCRαβ data evaluated on the same cross validation public test sets (Figure 2D; solid lines). Models trained on TCRs recognizing the A0201_GILGFVFTL epitope, which is the only SEQTR-profiled epitope also present in both the ePytope-TCR and IMMREP23 test sets, were further assessed on these independent test sets (Figure 2D; dash-dotted lines for ePytope-TCR and dashed lines for IMMREP23).

For A0101_SALPTNADLY, A0101_TDLGQNLLY and A0101_VSDGGPNLY, which were profiled by SEQTR in this study and are present in the IMMREP23 dataset, SEQTR-trained models were evaluated on the corresponding IMMREP23 test sets and compared to published pretrained versions of the tools (Figure 3B). Reported values (Figure 3B) represent the mean AUC01 per epitope. The TSP profiles (Figure S3) were plotted following the approach described in Liu *et al*. (see also https://tcrmotifatlas.unil.ch/Building_motifs) ^5^. In these TSP profiles, for the SEQTR data, a SEQTR baseline was used to account for expected differences between different TCR-sequencing protocols.

### AlphaFold3 predictions

Predictions for the three epitopes with little or no training data in IMMREP23 (A0101_SALPTNADLY, A0101_TDLGQNLLY and A0101_VSDGGPNLY) were generated using AlphaFold3, both with a local implementation of the tool and via the AlphaFold3 webserver. For each prediction, the five chains were modelled jointly: TCRα, TCRβ, the peptide, the MHC and the β2m. The top-ranked structure (rank 0, highest confidence) was always selected. The sum of the chain pair ipTM scores corresponding to the peptide/TCRα, peptide/TCRβ, MHC/TCRα and MHC/TCRβ interactions was extracted from the AlphaFold3 outputs and used to compute AUC01 values.

### Model performance evaluation packages

AUC and norsmalized AUC01 were computed using the roc() function from the pROC (v1.19.0.1) R package with parameters “levels = c(0, 1) and direction = <” ^43^. Paired Wilcoxon tests were performed using the wilcox_test() function from the rstatix (v0.7.2) R package ^44^.

## Contributions

DG designed the study and coordinated the project. AS analysed the data, compiled the final results and built the Figures. AA, RG performed the experiments. DM, YL, GC, JR contributed to the data analysis. AH supervised the experiments. DG wrote the manuscript, with input from all co-authors.s

## Acknowledgments

We are thankful to Julien Schmidt and the Peptide and Tetramer Core Facility for the synthesis of the peptides. The work was supported by the SNF Project Grant (320030-231333) and the ISREC foundation. “Figure 1A and Figure 3A” were created in BioRender (edited in inkscape). Shah, A. (https://BioRender.com/j4gtgel) is licensed under CC BY 4.0.

## References

1. Dash, P., Fiore-Gartland, A.J., Hertz, T., Wang, G.C., Sharma, S., Souquette, A., Crawford, J.C., Clemens, E.B., Nguyen, T.H.O., Kedzierska, K., et al. (2017). Quantifiable predictive features define epitope-specific T cell receptor repertoires. Nature 547, 89–93. 10.1038/nature22383.

2. Glanville, J., Huang, H., Nau, A., Hatton, O., Wagar, L.E., Rubelt, F., Ji, X., Han, A., Krams, S.M., Pettus, C., et al. (2017). Identifying specificity groups in the T cell receptor repertoire. Nature 547, 94–98. 10.1038/nature22976.

3. Jokinen, E., Huuhtanen, J., Mustjoki, S., Heinonen, M., and Lähdesmäki, H. (2021). Predicting recognition between T cell receptors and epitopes with TCRGP. PLoS Comput. Biol. 17, e1008814. 10.1371/journal.pcbi.1008814.

4. Springer, I., Tickotsky, N., and Louzoun, Y. (2021). Contribution of T Cell Receptor Alpha and Beta CDR3, MHC Typing, V and J Genes to Peptide Binding Prediction. Front. Immunol. 12. 10.3389/fimmu.2021.664514.

5. Liu, Y., Croce, G., Tadros, D., Moreno, D., Michel, A., Thierry, A.-C., Genolet, R., Perez, M.A., Lani, R., Guillaume, P., et al. (2025). Key determinants of T cell epitope recognition revealed by TCR specificity profiles. Preprint at bioRxiv, 10.1101/2025.11.17.688817.

6. Henderson, J., Nagano, Y., Milighetti, M., and Tiffeau-Mayer, A. (2024). Limits on inferring T cell specificity from partial information. Proc. Natl. Acad. Sci. 121, e2408696121. 10.1073/pnas.2408696121.

7. Robins, H.S., Campregher, P.V., Srivastava, S.K., Wacher, A., Turtle, C.J., Kahsai, O., Riddell, S.R., Warren, E.H., and Carlson, C.S. (2009). Comprehensive assessment of T-cell receptor beta-chain diversity in alphabeta T cells. Blood 114, 4099–4107. 10.1182/blood-2009-04-217604.

8. Heather, J.M., Ismail, M., Oakes, T., and Chain, B. (2018). High-throughput sequencing of the T-cell receptor repertoire: pitfalls and opportunities. Brief. Bioinform. 19, 554–565. 10.1093/bib/bbw138.

9. Genolet, R., Bobisse, S., Chiffelle, J., Arnaud, M., Petremand, R., Queiroz, L., Michel, A., Reichenbach, P., Cesbron, J., Auger, A., et al. (2023). TCR sequencing and cloning methods for repertoire analysis and isolation of tumor-reactive TCRs. Cell Rep. Methods 3. 10.1016/j.crmeth.2023.100459.

10. Wang, C., Sanders, C.M., Yang, Q., Schroeder, H.W., Wang, E., Babrzadeh, F., Gharizadeh, B., Myers, R.M., Hudson, J.R., Davis, R.W., et al. (2010). High throughput sequencing reveals a complex pattern of dynamic interrelationships among human T cell subsets. Proc. Natl. Acad. Sci. U. S. A. 107, 1518–1523. 10.1073/pnas.0913939107.

11. Rosati, E., Dowds, C.M., Liaskou, E., Henriksen, E.K.K., Karlsen, T.H., and Franke, A. (2017). Overview of methodologies for T-cell receptor repertoire analysis. BMC Biotechnol. 17, 61. 10.1186/s12896-017-0379-9.

12. Lee, E.S., Thomas, P.G., Mold, J.E., and Yates, A.J. (2017). Identifying T Cell Receptors from High-Throughput Sequencing: Dealing with Promiscuity in TCRα and TCRβ Pairing. PLoS Comput. Biol. 13, e1005313. 10.1371/journal.pcbi.1005313.

13. Holec, P.V., Berleant, J., Bathe, M., and Birnbaum, M.E. (2019). A Bayesian framework for high-throughput T cell receptor pairing. Bioinformatics 35, 1318–1325. 10.1093/bioinformatics/bty801.

14. Stubbington, M.J.T., Lönnberg, T., Proserpio, V., Clare, S., Speak, A.O., Dougan, G., and Teichmann, S.A. (2016). T cell fate and clonality inference from single cell transcriptomes. Nat. Methods 13, 329–332. 10.1038/nmeth.3800.

15. Pai, J.A., and Satpathy, A.T. (2021). High-throughput and single-cell T cell receptor sequencing technologies. Nat. Methods 18, 881–892. 10.1038/s41592-021-01201-8.

16. Howie, B., Sherwood, A.M., Berkebile, A.D., Berka, J., Emerson, R.O., Williamson, D.W., Kirsch, I., Vignali, M., Rieder, M.J., Carlson, C.S., et al. (2015). High-throughput pairing of T cell receptor α and β sequences. Sci. Transl. Med. 7, 301ra131–301ra131. 10.1126/scitranslmed.aac5624.

17. Pogorelyy, M.V., Kirk, A.M., Adhikari, S., Minervina, A.A., Sundararaman, B., Vegesana, K., Brice, D.C., Scott, Z.B., and Thomas, P.G. (2026). TIRTL-seq: deep, quantitative and affordable paired TCR repertoire sequencing. Nat. Methods 23, 56–64. 10.1038/s41592-025-02907-9.

18. Mayer-Blackwell, K., Schattgen, S., Cohen-Lavi, L., Crawford, J.C., Souquette, A., Gaevert, J.A., Hertz, T., Thomas, P.G., Bradley, P., and Fiore-Gartland, A. (2021). TCR meta-clonotypes for biomarker discovery with tcrdist3 enabled identification of public, HLA-restricted clusters of SARS-CoV-2 TCRs. eLife 10, e68605. 10.7554/eLife.68605.

19. Weber, A., Born, J., and Rodriguez Martínez, M. (2021). TITAN: T-cell receptor specificity prediction with bimodal attention networks. Bioinformatics 37, i237–i244. 10.1093/bioinformatics/btab294.

20. Croce, G., Bobisse, S., Moreno, D.L., Schmidt, J., Guillame, P., Harari, A., and Gfeller, D. (2024). Deep learning predictions of TCR-epitope interactions reveal epitope-specific chains in dual alpha T cells. Nat. Commun. 15, 3211. 10.1038/s41467-024-47461-8.

21. Jensen, M.F., and Nielsen, M. (2024). NetTCR 2.2 - Improved TCR specificity predictions by combining pan- and peptide-specific training strategies, loss-scaling and integration of sequence similarity. eLife 12. 10.7554/eLife.93934.2.

22. Meynard-Piganeau, B., Feinauer, C., Weigt, M., Walczak, A.M., and Mora, T. (2024). TULIP: A transformer-based unsupervised language model for interacting peptides and T cell receptors that generalizes to unseen epitopes. Proc. Natl. Acad. Sci. 121, e2316401121. 10.1073/pnas.2316401121.

23. Nielsen, M., Eugster, A., Jensen, M.F., Goel, M., Tiffeau-Mayer, A., Pelissier, A., Valkiers, S., Martínez, M.R., Meynard-Piganeeau, B., Greiff, V., et al. (2024). Lessons learned from the IMMREP23 TCR-epitope prediction challenge. ImmunoInformatics 16, 100045. 10.1016/j.immuno.2024.100045.

24. Drost, F., Chernysheva, A., Albahah, M., Kocher, K., Schober, K., and Schubert, B. (2025). Benchmarking of T cell receptor-epitope predictors with ePytope-TCR. Cell Genomics, 100946. 10.1016/j.xgen.2025.100946.

25. Sidhom, J.-W., Larman, H.B., Pardoll, D.M., and Baras, A.S. (2021). DeepTCR is a deep learning framework for revealing sequence concepts within T-cell repertoires. Nat. Commun. 12, 1605. 10.1038/s41467-021-21879-w.

26. Zhang, W., Hawkins, P.G., He, J., Gupta, N.T., Liu, J., Choonoo, G., Jeong, S.W., Chen, C.R., Dhanik, A., Dillon, M., et al. (2021). A framework for highly multiplexed dextramer mapping and prediction of T cell receptor sequences to antigen specificity. Sci. Adv. 7, eabf5835. 10.1126/sciadv.abf5835.

27. Kim, H.Y., Kim, S., Park, W.-Y., and Kim, D. (2023). TSpred: a robust prediction framework for TCR-epitope interactions based on an ensemble deep learning approach using paired chain TCR sequence data. Preprint, 10.1101/2023.12.04.570002.

28. Korpela, D., Jokinen, E., Dumitrescu, A., Huuhtanen, J., Mustjoki, S., and Lähdesmäki, H. (2023). EPIC-TRACE: predicting TCR binding to unseen epitopes using attention and contextualized embeddings. Bioinforma. Oxf. Engl. 39, btad743. 10.1093/bioinformatics/btad743.

29. Abramson, J., Adler, J., Dunger, J., Evans, R., Green, T., Pritzel, A., Ronneberger, O., Willmore, L., Ballard, A.J., Bambrick, J., et al. (2024). Accurate structure prediction of biomolecular interactions with AlphaFold 3. Nature 630, 493–500. 10.1038/s41586-024-07487-w.

30. Boitreaud, J., Discovery, C., Dent, J., McPartlon, M., Meier, J., Reis, V., Rogozhnikov, A., and Wu, K. (2024). Chai-1: Decoding the molecular interactions of life. Preprint at bioRxiv, 10.1101/2024.10.10.615955.

31. Passaro, S., Corso, G., Wohlwend, J., Reveiz, M., Thaler, S., Somnath, V.R., Getz, N., Portnoi, T., Roy, J., Stark, H., et al. (2025). Boltz-2: Towards Accurate and Efficient Binding Affinity Prediction. BioRxiv Prepr. Serv. Biol., 2025.06.14.659707. 10.1101/2025.06.14.659707.

32. Bradley, P. (2023). Structure-based prediction of T cell receptor:peptide-MHC interactions. eLife 12, e82813. 10.7554/eLife.82813.

33. Deleuran, S.N., and Nielsen, M. (2025). NetTCR-struc, a structure driven approach for prediction of TCR-pMHC interactions. Front. Immunol. 16. 10.3389/fimmu.2025.1616328.

34. Messemaker, M., Kwee, B.P.Y., Moravec, Ž., Álvarez-Salmoral, D., Urbanus, J., de Paauw, S., Geerligs, J., Voogd, R., Morris, B., Guislain, A., et al. (2025). A functionally validated TCR-pMHC database for TCR specificity model development. bioRxiv, 2025.04.28.651095. 10.1101/2025.04.28.651095.

35. Goncharov, M., Bagaev, D., Shcherbinin, D., Zvyagin, I., Bolotin, D., Thomas, P.G., Minervina, A.A., Pogorelyy, M.V., Ladell, K., McLaren, J.E., et al. (2022). VDJdb in the pandemic era: a compendium of T cell receptors specific for SARS-CoV-2. Nat. Methods 19, 1017–1019. 10.1038/s41592-022-01578-0.

36. Vita, R., Blazeska, N., Marrama, D., IEDB Curation Team Members, Duesing, S., Bennett, J., Greenbaum, J., De Almeida Mendes, M., Mahita, J., Wheeler, D.K., et al. (2025). The Immune Epitope Database (IEDB): 2024 update. Nucleic Acids Res. 53, D436–D443. 10.1093/nar/gkae1092.

37. Shugay, M., Bagaev, D.V., Zvyagin, I.V., Vroomans, R.M., Crawford, J.C., Dolton, G., Komech, E.A., Sycheva, A.L., Koneva, A.E., Egorov, E.S., et al. (2018). VDJdb: a curated database of T-cell receptor sequences with known antigen specificity. Nucleic Acids Res. 46, D419–D427. 10.1093/nar/gkx760.

38. 10x Genomics (2019). A New Way of Exploring Immunity - Linking Highly Multiplexed Antigen Recognition to Immune Repertoire and Phenotype. Immunol. Microbiol. Technol. Netw. http://www.technologynetworks.com/immunology/application-notes/a-new-way-of-exploring-immunity-linking-highly-multiplexed-antigen-recognition-to-immune-repertoire-332554.

39. Tickotsky, N., Sagiv, T., Prilusky, J., Shifrut, E., and Friedman, N. (2017). McPAS-TCR: a manually curated catalogue of pathology-associated T cell receptor sequences. Bioinformatics 33, 2924–2929. 10.1093/bioinformatics/btx286.

40. Moreno, D.L., Croce, G., and Gfeller, D. (2025). Statistical modelling of CDR3 sequences provides robust quality control for TCR repertoire datasets. Preprint at bioRxiv, 10.64898/2025.12.11.693618.

41. Schmidt, F., Fields, H.F., Purwanti, Y., Milojkovic, A., Salim, S., Wu, K.X., Simoni, Y., Vitiello, A., MacLeod, D.T., Nardin, A., et al. (2023). In-depth analysis of human virus-specific CD8+ T cells delineates unique phenotypic signatures for T cell specificity prediction. Cell Rep. 42, 113250. 10.1016/j.celrep.2023.113250.

42. Bolotin, D.A., Poslavsky, S., Mitrophanov, I., Shugay, M., Mamedov, I.Z., Putintseva, E.V., and Chudakov, D.M. (2015). MiXCR: software for comprehensive adaptive immunity profiling. Nat. Methods 12, 380–381. 10.1038/nmeth.3364.

43. Robin, X., Turck, N., Hainard, A., Tiberti, N., Lisacek, F., Sanchez, J.-C., and Müller, M. (2011). pROC: an open-source package for R and S+ to analyze and compare ROC curves. BMC Bioinformatics 12, 77. 10.1186/1471-2105-12-77.

44. Kassambara, A. (2023). rstatix: Pipe-Friendly Framework for Basic Statistical Tests. Version 0.7.3.

