## Supplementary Figures and Supplementary Table Captions for "Unpaired TCRα + TCRβ sequencing is sufficient for training machine learning TCR-epitope recognition predictors"

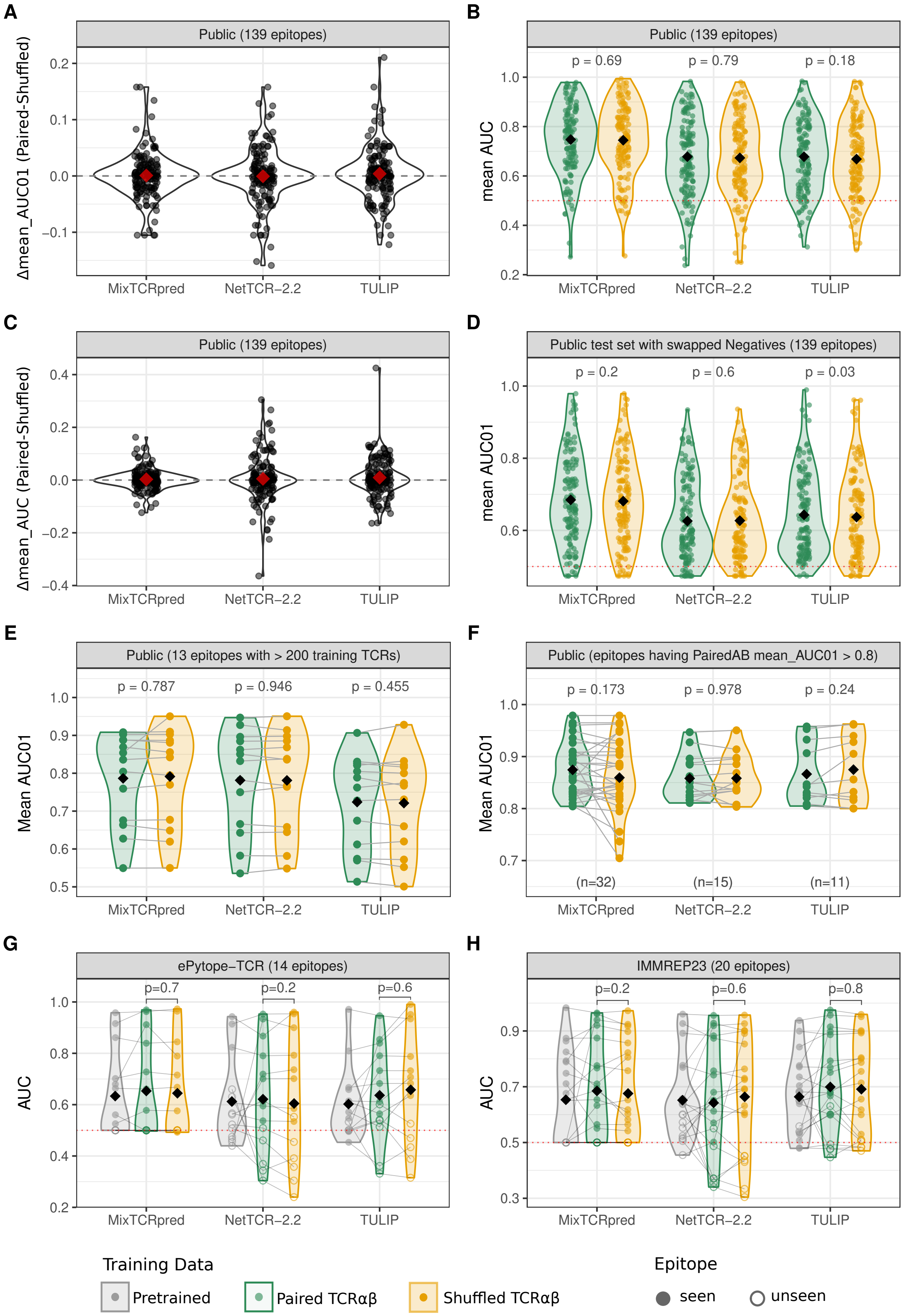

**Figure S1: Shuffling TCRαβ pairs used for training TCR-epitope interaction predictors preserves their accuracy. A)** Difference between mean AUC01 values of 5 cross-validation test sets obtained when training different tools on paired TCRαβ or shuffled TCRαβ sequences. **B)** Mean AUC values of 5 cross-validation test sets obtained when training different tools on paired TCRαβ or shuffled TCRαβ sequences. **C)** Difference between mean AUC values shown in S1B. **D)** Cross-validation AUC01 values obtained when training different tools on paired TCRαβ or shuffled TCRαβ sequences and using swapped negatives in the test sets. **E)** Cross-validation AUC01 values obtained when training different tools on paired TCRαβ or shuffled TCRαβ sequences for epitopes having at least 200 TCRs in training data. **F)** Cross-validation AUC01 values of epitopes having AUC01>=0.8 when training different tools on paired TCRαβ. **G)** AUC values for epitopes in the ePytope-TCR benchmark when training different tools on paired TCRαβ or shuffled TCRαβ sequences. **H)** AUC values for epitopes in the IMMREP23 benchmark when training different tools on paired TCRαβ or shuffled TCRαβ sequences. P-values were computed using the Wilcoxon paired test. Black diamonds show the mean values across all epitopes. For comparison, AUC obtained with the published models (i.e., without retraining) of each tool are shown in grey in panels G-H.

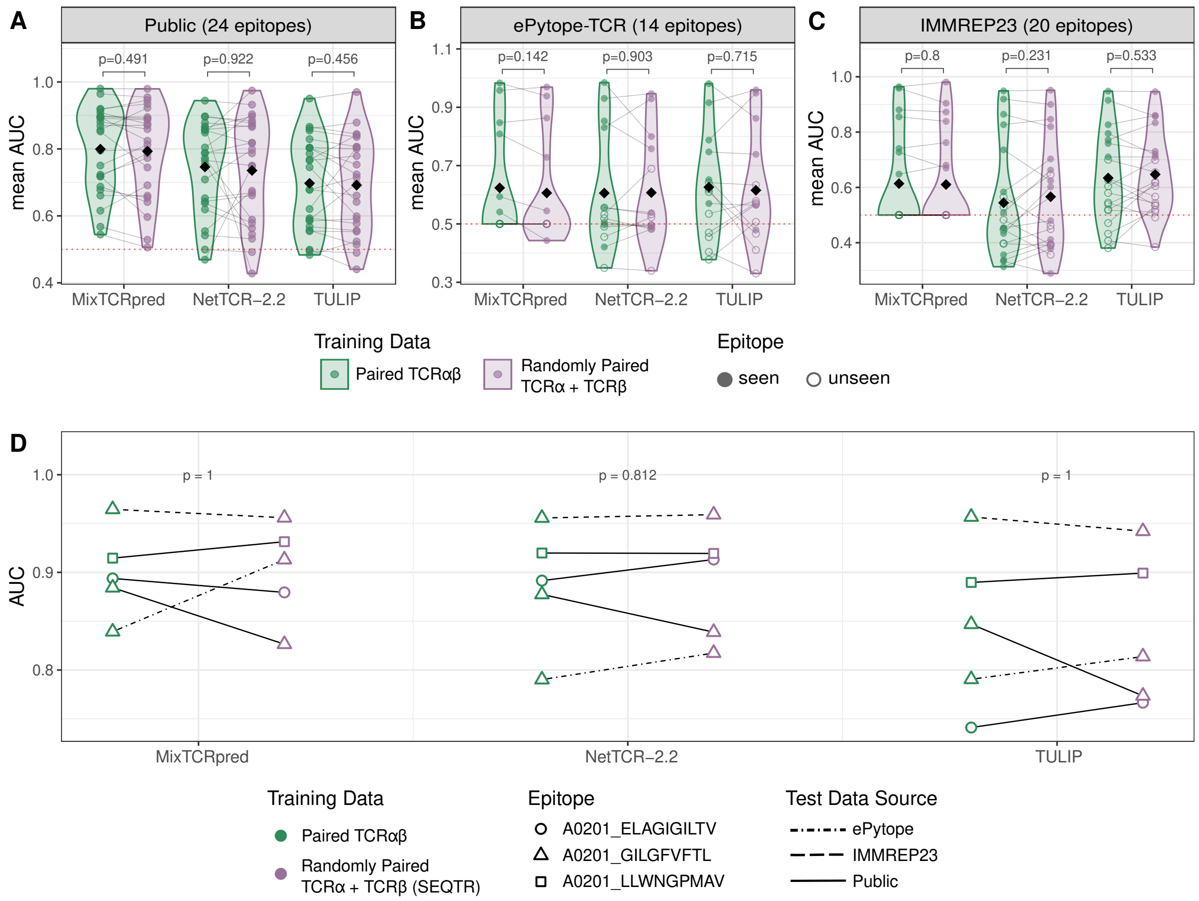


**Figure S2: Unpaired TCRα + TCRβ sequences can be used for training TCR-epitope interaction predictors. A)** Cross-validation AUC values obtained when training tools on paired TCRαβ or unpaired TCRα + TCRβ data. **B)** AUC values for epitopes in the ePytope-TCR benchmark when training tools on paired TCRαβ or unpaired TCRα + TCRβ data. **C)** AUC values for epitopes in the IMMPEP23 benchmark when training tools on paired TCRαβ or unpaired TCRα + TCRβ data. **D)** AUC values for three epitopes (GILGFVFTL, LLWNGPMAV and ELAGIGILTV, all restricted to HLA-A*02:01) when training tools on paired TCRαβ or newly generated unpaired TCRα + TCRβ data with the SEQTR protocol. GILGFVFTL predictions were evaluated on three test sets indicated by different line types. P-values were computed with the Wilcoxon paired test.

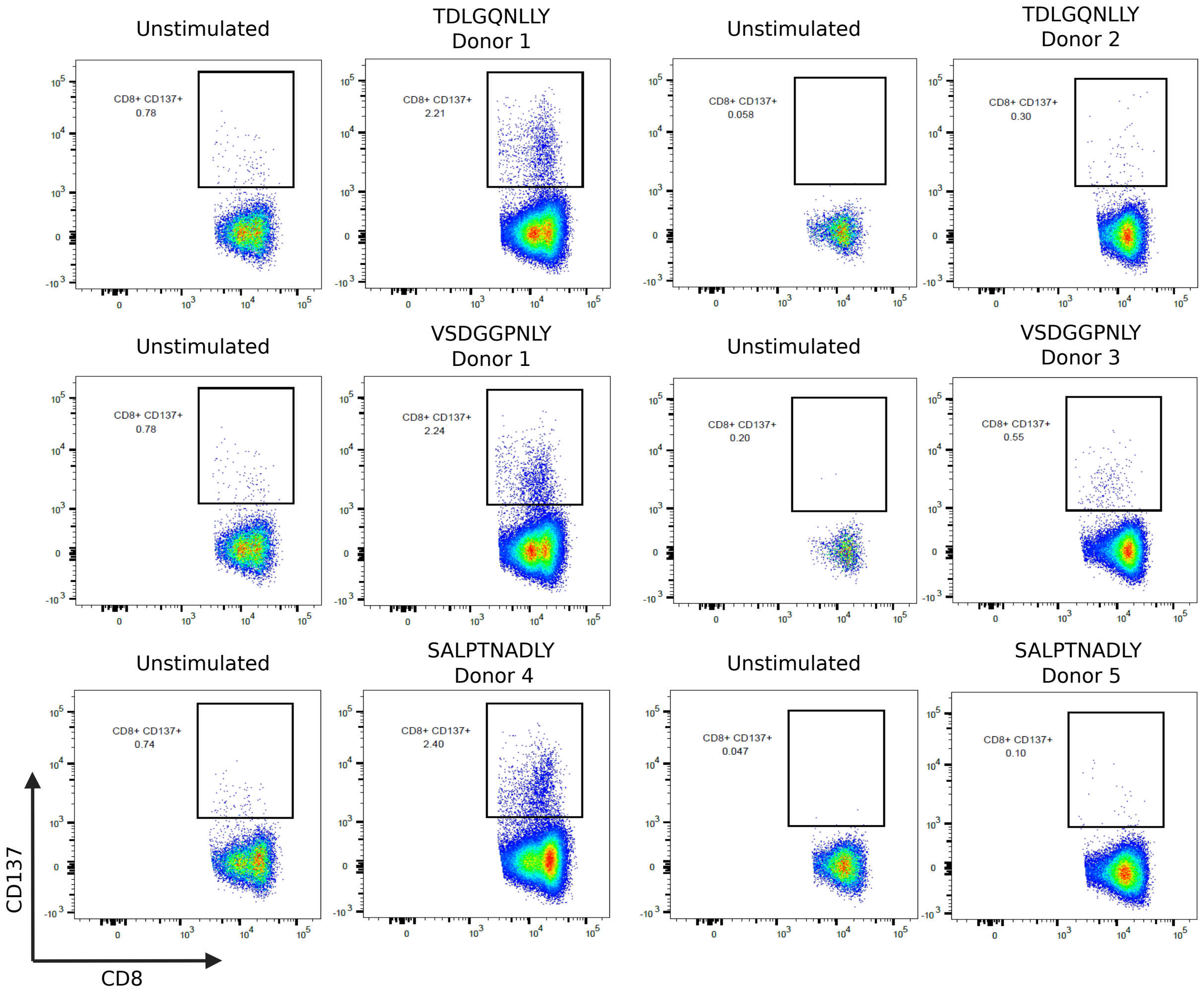


**Figure S3:** Flow cytometry (FACS) plots for the isolation of CD8 T cells based on CD137 expression after stimulation with the TDLGQNLLY, VSDGGPNLY, and SALPTNADLY peptides. Negative controls correspond to unstimulated samples. Data is shown for two independent donors per peptide.

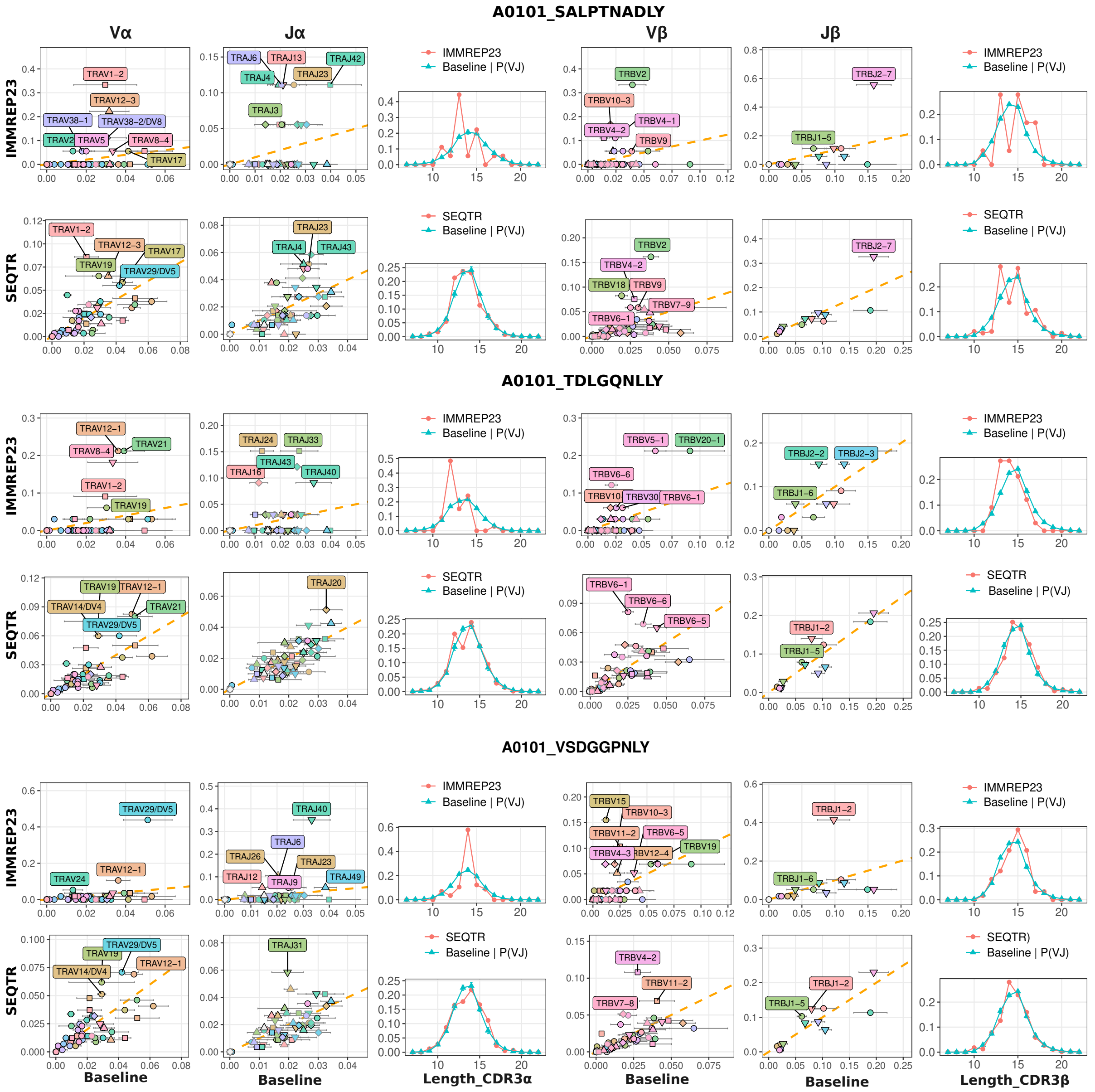


**Figure S4:** Comparison between the TCR specificity profiles based on the IMMREP23 solution and the newly generated SEQTR data for three epitopes with very limited or no publicly available training data at the time of the IMMREP23 competition. For IMMREP23 data, the default baseline was used in the TCR specificity profiles. For SEQTR data, a dedicated baseline derived from bulk repertoires sequenced with SEQTR was used.

### Supplementary Tables

**Table S1:** Training and test data used in the cross-validation analysis to compare tools trained on paired TCRαβ and randomly shuffled TCRαβ.

**Table S2:** Training and test data used in the cross-validation analysis to compare tools trained on paired TCRαβ or randomly paired TCRα + TCRβ.

**Table S3:** Newly generated SEQTR data for three epitopes in IMMREP23 (A0101_SALPTNADLY, A0101_TDLGQNLLY and A0101_ VSDGGPNLY).

**Table S4:** Training data used to compare Randomly Paired (using 5 different seeds) SEQTR TCRα + TCRβ with pretrained models for three epitopes in IMMREP23 (A0101_SALPTNADLY, A0101_TDLGQNLLY and A0101_ VSDGGPNLY).
